# The nascent RNA labelling compound 5-ethynyl uridine (EU) integrates into DNA in some animals

**DOI:** 10.1101/2024.10.22.619599

**Authors:** Malin A. Kjosavik, Katherine L. P. Downham, Ruth Styfhals, Leonie Adelmann, Marios Chatzigeorgiou, Florian Raible, Pawel Burkhardt, Fergal O’Farrell, Patrick R. H. Steinmetz, Kathrin Garschall

## Abstract

**Background:** The detection of *de novo* synthesized mRNA transcripts is crucial for understanding the regulation of eukaryotic transcription. Using nucleoside or nucleotide analogues to label nascent RNA is potentially jeopardized by the ubiquitous presence of ribonucleotide reductase enzymes (RNRs) that can convert ribonucleotides into 2’-deoxyribonucleotides, the building blocks of DNA. Despite this challenge, the uridine analogue 5-ethynyl uridine (EU) has been commercialized and routinely used as specific label for nascent RNAs. Here, we employ confocal imaging, flow cytometry and biochemistry methods to study the specificity of EU to label RNA in six different animal species.

**Results:** We demonstrate that EU integrates as expected predominantly into RNA of human embryonic kidney cell line (HEK293), the *Drosophila* wing disc and the comb jelly *Mnemiopsis leidyi*. In contrast, we found that EU predominantly labels DNA in the sea anemones *Nematostella vectensis* and *Exaiptasia diaphana*, and the polychaete *Platynereis dumerilii*. In *Nematostella*, we show that inhibiting RNR by hydroxyurea abolishes cell proliferation and the incorporation of EU into DNA. Alternative compounds for labelling nascent RNA, such as 5-ethynyl cytidine (EC), 5-ethynyl uridine triphosphate (EUTP) or 2-ethynyl adenosine (EA) show similarly low specificity for RNA in *Nematostella*.

**Conclusions:** Our findings raise concerns about the specificity of ethynylated nucleosides and nucleotides, including EU, to label RNA in some animals. We therefore suggest good practice guidelines for using EU as an RNA labelling tool and discuss pitfalls and indicators that help identifying unintentional DNA labelling.

## Background

The regulation of eukaryotic gene expression is crucial for maintaining basic cellular functions and adapting to environmental changes. In multicellular organisms (e.g., animals, plants), transcriptional regulation ensures that genes are expressed at the right time, place and levels to control embryonic development and adult homeostasis [1-3]. Intricate gene regulatory networks underlie the development and evolution of complex cell types and cellular specializations in animals [1, 4, 5], plants [6] and some unicellular eukaryotes such as *Plasmodium sp.* [7]. A key aspect of studying gene regulation is the detection, quantification and analysis of changes in *de novo* mRNA synthesis. As standard RNA extraction and detection procedures cannot discriminate between stored and actively transcribed mRNAs, modified nucleosides (i.e. nucleobase bound to a ribose) have been introduced in the 1960s to label and isolate nascent RNAs [8-11]. Nucleosides need to be tri-phosphorylated by specific nucleoside kinases to nucleoside triphosphates (NTPs) before they can be used as substrate by RNA polymerases to synthesise RNAs [12]. The intermediate products of NTPs, the nucleoside diphosphates (NDPs), are however not only a precursor of NTPs and RNA synthesis. Instead, the ribonucleotide reductase (RNR) enzymes reduce NDPs to 2’-deoxy-NDPs (dNDPs) that can then act as precursors for DNA synthesis [12]. This function, discovered decades ago, is an essential and rate-limiting step for DNA replication that is highly conserved across bacteria, eukaryotes and archaea [13-15]. Consequently, the use of modified nucleotides carries a risk of not only labelling nascent RNA but also replicating DNA. Indeed, radioactively labelled nucleosides developed decades ago to quantify nascent RNA synthesis via autoradiography (e.g., ^3^H-uridine, ^3^H-cytidine or the uracil precursor ^3^H-orotic acid) have been repeatedly found to integrate at significant levels into DNA [8, 16]. Furthermore, the pyrimidine analogue 5-fluorouracil (5-FU), used in chemotherapy, was detected in both RNA (as 5-FU triphosphate; 5-FUTP) and DNA (as 5-fluoro-2’-deoxyuridine-5’-triphosphate; 5-FdUTP) of cancer cells [17-19]. Similarly, the ribonucleosides cytosine arabinoside (cytarabine or Ara-C) and 9-β-D-arabinofuranosyl-2-fluoroadenine (F-ara-A) integrate into both DNA and RNA primers during DNA synthesis [20]. The resulting inhibition of DNA replication has been exploited in their development to chemotherapeutics [20, 21].

The more recently developed nucleoside 5-ethynyl uridine (EU) can be covalently coupled to azide-containing fluorophores through ‘click chemistry’ in a one-step, copper-catalysed reaction [22]. This approach avoids harsh denaturation steps necessary for immunodetection and offers enhanced safety and detection sensitivity. EU was reported to specifically label RNA in mammalian cell lines (NIH 3T3) and was suggested to constitute a poor substrate for RNR [22]. Since its introduction in 2008, EU has been widely adopted to visualise and quantify global nascent RNA synthesis and turnover in eukaryotic cells or tissues using confocal microscopy. Examples include plants [23] and various animals such as mice [22], *Drosophila* [24], snails [25, 26] and cnidarians such as *Hydractinia* [27] or *Nematostella* [28]. However, the possibility of dual integration into both RNA and DNA is not always explicitly considered when EU is used for nascent RNA labelling.

Here, we aimed to validate the use of EU as a tool for nascent RNA labelling by visualising and quantifying *de novo* transcriptional levels in the sea anemone *Nematostella vectensis.* Against our expectations, EU and several other ethynylated nucleosides or nucleotides intensely label replicating DNA of proliferative cells in addition to RNA. We therefore expanded our evaluation of EU as nascent RNA labelling tool to other animal systems, including biomedically relevant models. We confirmed that EU predominantly incorporates into RNA in the human cell line HEK293T, the wing disc in flies (*Drosophila melanogaster*) and comb jellies (*Mnemiopsis* leidyi). Strikingly, however, EU predominantly labelled DNA also in the sea anemone *Exaiptasia diaphana* and the polychaete annelid *Platynereis dumerlii*. The consistent dual labelling of DNA and RNA renders EU therefore unsuitable for visualising and quantifying *de novo* RNA transcripts in *Nematostella,* Aiptasia and *Platynereis*. In consequence, we critically evaluate the use of EU as RNA labelling tool in the previous literature and propose guidelines to prevent mis-identifying replicating DNA as nascent RNA.

## Methods

### Animal culture *Nematostella*

Adult *Nematostella vectensis* polyps derive from the original CH6 female and CH2 male lab strains and were cultured as previously described [29, 30]. Shortly, adults were kept at 18°C in 16‰ salt concentration. Juvenile *Nematostella vectensis* polyps result from crosses between wild type animals and were raised at 25°C in a darkened incubator at 16‰ salt concentration (*Nematostella* medium). Polyps were fed 5x/week with *Artemia* nauplii and grown until the second pair of mesenteries had developed, when they were defined as ‘juveniles’. The culture medium was replaced on the same day after each feeding.

### EU, EdU, EC, EUTP or EA labelling and co-labelling with BrdU

Working concentrations for EU (5-ethynyl uridine, Invitrogen) were determined by testing the fraction and fluorescent signal intensity of EU-labelled cells (EU index) by flow cytometry after incubation of 0.5mM, 1mM and 5mM. As the EU index was similar between 0.5mM and 1mM but reduced in the 5mM EU treatment (Supplementary Fig. 2A, Additional file 1) all labelling experiments were carried out at 0.5mM EU.

*Nematostella* juveniles were prepared for incubations by relaxing them for 15 min in 50mM MgCl_2_ in *Nematostella* medium. Relaxed animals were then transferred to labelling medium containing 2% DMSO, 50mM MgCl_2_ and either 0.5mM EU, 0.5mM EdU (5-ethynyl 2’-deoxyuridine, Invitrogen), 0.5mM EC (5-ethynyl cytidine, Jena Bioscience), 0.5mM EUTP (5-ethynyl UTP, Jena Bioscience) or 0.5mM EA (2-ethynyl adenosine, Jena Bioscience). In co-labelling assays, 0.1mM BrdU (5-Bromo-2’-deoxyuridine, Invitrogen) was combined with 0.5mM EU/EdU in *Nematostella* medium with 2% DMSO and 50mM MgCl_2_.

All incubations were carried out for 2h at RT. For unlabelled controls, animals were exposed to 2% DMSO and 50mM MgCl_2_ in culture medium. Up to 15 animals were incubated per single well of a 12-well plate, or up to 50 animals per well of a 6-well plate. Following incorporation, the animals were rinsed with *Nematostella* medium containing 50mM MgCl_2_ before fixation and dissociation.

### Fixation and sample preparation for whole mount confocal microscopy

After labelling with EU, EdU, EC, EUTP, EA and/or BrdU, the relaxed polyps were transferred gently into fixation solution (1x PBS, 3.7% FA, 0.2% TritonX, 0.5% DMSO), incubated for 30 min at RT followed by an over-night fixation in fresh fixative at 4°C. After fixation, the polyps were washed 3-5x with PTx (1x PBS, 0.2% TritonX) for permeabilization, briefly blocked in PBTx (1x PBS, 0.2% TritonX, 3% BSA) and rinsed with 1x PBS. The Click-iT™ reaction cocktail containing an Alexa fluorophore azide (Alexa Fluor™ 488 or 555) was added to all samples, including unlabelled controls, and incubated for 30 min at RT in the dark according to manufacturer protocol (Click-iT™ Nascent RNA Capture Kit /cell proliferation kit, Invitrogen). Finally, the polyps were washed 3-5x with PTx (1x PBS, 0.2% TritonX), incubated with Hoechst33342 (1:5000) and mounted for confocal microscopy.

### Cell maceration using ACME and Click-iT™ reaction on cell suspensions

For blotting extracted DNA or RNA as well as to quantify the proportion and signal intensity of EU+ cells by flow cytometry, polyps were dissociated and fixed after labelling by adapting the ‘<u>AC</u>etic acid/<u>ME</u>thanol’ (ACME) maceration protocol to *Nematostella* juveniles as previously described [31, 32]. In short, 5ml of fresh ACME solution (15% methanol/10% glacial acetic acid/10% glycerol in Milli-Q water) were added to labelled polyps in gentleMACS™ C-tubes (Miltenyi Biotec) and incubated for 1 hour at RT on a see-saw shaker. During incubation, the tissue was disrupted using a gentleMACS™ tissue homogenizer (Miltenyi Biotec) by running the program “B_01” for initial dissociation after 10 min ACME incubation, followed by four runs of the “Multi_A_01” programme over the following 50 min until all tissue was dissociated. The resulting cell suspension was washed twice with 10ml of ice-cold 1×PBS/1% BSA, pelleted for 5min at 800*g* at 4°C, resuspended in 1ml of 1× PBS/1% BSA and filtered through a pre-wetted 50µm CellTrics strainer (Sysmex). The resulting cells were permeabilized with PTx (1x PBS, 0.2% TritonX) at RT for 15 min, washed with 1x PBS and incubated for 30 min at RT in the dark with a modified Click reaction cocktail (Click-iT™ Nascent RNA kit, Invitrogen) where the reaction buffer was replaced by 1x PBS. The cells were then pelleted by spinning for 5min at 800*g* at 4°C, washed with PTx and transferred to 1xPBS containing 1% BSA for short term storage at 4°C. For long term storage, cells were resuspended in 90% Methanol and kept at -20°C.

### Detection of BrdU and RNA Pol II-Ser2 immunofluorescence

In samples that were co-labelled with BrdU, the incorporated EU/EdU was Click-ed to Alexa Fluor™ 555 (Invitrogen) prior to BrdU immunofluorescence detection. The tissue was first incubated with 2M HCl in 1xPBS for 30 min at RT to make the DNA accessible, followed by multiple washes with PBTx (1x PBS, 3% BSA, 0.2% TritonX) and a blocking step for 2h at RT in blocking solution (5% NGS, 3% BSA and 0.02% TritonX in 1x PBS). The primary anti-BrdU rabbit antibody (Invitrogen, PA5-32256) was 1:500 diluted in blocking solution and incubated o/n at 4°C on a rotation shaker without inversion. After 3 washes with PBTx, the tissue was blocked for 1h at RT in freshly prepared blocking solution and incubated for 2h at RT with 1:1000 goat-anti-rabbit-Alexa488 antibody (Invitrogen, A-11034), 1:5000 Hoechst33342 (Life Technologies) in blocking solution. Samples were then washed 3 times in PTx to reduce unspecific signal.

For the immunofluorescence labelling of RNA Pol II CTD repeat YSPTSPS phospho-Ser2 (AB5095, Abcam), indicative of active transcription, fixed, click-ed polyps were washed and blocked for 1h in blocking solution (5% NGS, 3% BSA and 0.02% TritonX in 1x PBS). Primary antibody (AB5095, Abcam) was diluted 1:500 in blocking solution and incubated at 4°C over night on a rotation shaker without inversion. After 5 washes with PBTx (1x PBS, 3% BSA, 0.2% TritonX) and blocking for 30 min in blocking solution at RT, secondary goat-anti-rabbit-Alexa488 antibody (Invitrogen, A-11034) was added 1:1000 along with 1:5000 Hoechst33342 and kept for 2h at RT. Samples were then washed in PTx for 3 times and mounted for confocal microscopy.

### Confocal imaging

For whole mount images, fixed and stained polyps were cut transversally using a razorblade, and mid-section pieces were mounted on slides using ProLong Gold^TM^ (Invitrogen). Confocal imaging was performed using an Olympus FLUOVIEW FV3000 microscope with standard PMT detectors and a 60x silicon-immersion lens. Laser power and sensor sensitivity settings were kept identical between acquisitions within each of these groups of images: (i) strong nuclear EU label in proliferating cells (Fig. 2A, C, E, G, I, K); (iia) nucleolar EU/EdU label (Fig. 4A-D’’) or (iib) Fig. 4E-H; (iii) EdU label (Fig. 2B, D, F, H, J, L); (iv) EUTP nuclear label (Fig. 5A-B); (v) EUTP nucleolar label (Fig. 5A’-B’); (vi) EC nuclear label (Fig. 5C-D); (vii) EC nucleolar label (Fig. 5C’-D’); (viii) EA label (Fig. 5E-F); (ix) EU/BrdU label in HEK293T cells (Fig. 6F-G’’’); (x) EU/BrdU label in *D. melanogaster* (Fig. 6H-I’’’); (xi) EU/BrdU label in *M. leidyi* (Fig. 6J-K’’’); (xii) EU label (Fig. 6L-L’,N-N’) and EdU label (Fig. 6M-M’,O-O’) in *P. dumerilii*; EdU/BrdU label in *E. diaphana* (Fig. 6P-Q’’’).

Images were consistently adjusted for brightness and contrast over all images within each group of images using the QuickFigures PlugIn [33] in Fiji/ImageJ [34, 35], except for the DNA stain that has been adjusted differently between images.

### Flow cytometry

For each of the experimental conditions (0.5mM, 1mM or 5mM EU), two (2% DMSO control) or three (all others) replicate pools of 7 juveniles each were dissociated and fixed using ACME and coupled to Alexa Fluor™ 555 as described above. The cell suspensions were stained with 1µg/ml FxCycle violet DNA dye (Invitrogen) at 4°C over-night and run on an LSRFortessa™ (BD Life Sciences) flow cytometer. EU-labelled cells were analysed using FlowJo version 10.8 (BD Life Sciences) (see gating strategy Supplementary Fig. 1, Additional file 1). Shortly, events were gated using FSC and SSC parameters before selecting for cells with a FxCycle violet DNA signal ranging between 2N and 4N. Within this cell population, EU positive cells were gated based on their Alexa555 signal above a threshold from unlabelled, 2% DMSO samples.

Individual cells were visualised after flow cytometry by adding 2 µg/ml Concanavalin A conjugated to Alexa Fluor™ 488 to label cytosol in EU-labelled cell suspensions.

### Flow cytometry data analysis

The fraction of EU positive cells out of all cells that show a cell cycle distribution based on DNA signal and their fluorescent intensity, was analysed between different concentration of EU (0.5mM/1mM/5mM) using One-way ANOVA. Pairwise comparisons between the different EU concentrations were calculated in Tukey’s multiple comparison test and adjusted p-values indicated as asterisks (Supplementary Fig. 2, Additional file 1).

### Hydroxyurea, actinomycin D and dNTPs incubations

For interventional treatments with 20µM hydroxyurea (Sigma, H8627) or 0.2µM or 8µM actinomycin D (Invitrogen, A7592), juveniles were pre-incubated for 30 min in the compound with 1% DMSO and 50mM MgCl_2_ in *Nematostella* medium. Subsequently, EU, EdU or DMSO was added into the medium to reach a final concentration of 0.5mM EU or EdU and 2% DMSO and polyps were incubated for 2 hours. For the treatment with 2mM dNTPs (New England Biolabs), 0.5mM per nucleotide to were added directly into the EU labelling medium containing 2% DMSO for a 2h incubation.

### Incubation in RNase A or RNase H

Following Click-iT^TM^ reaction, the samples were treated with 200µg/ml RNase A (PureLink™, Invitrogen) in PTx (0.2% TritonX, 1x PBS) for 15 min at 37°C, or 100U/ml RNase H (Thermo Fisher Scientific) in 1x RNase H Reaction Buffer for 30 min at 37°C. After incubation, the tissue was washed 3-5 times with PTx before mounting and confocal microscopy.

### Total RNA extraction for dot blot visualisation

Per sample, 50 EU-labelled animals were homogenized in 10ml of ACME solution, washed as described above but without filtering through CellTrics™ strainers. After Click reaction to Alexa Fluor™ 488, half of the resulting cell pellet was used for total RNA and half for genomic DNA extraction. For total RNA extraction, the cells were mixed with 1ml of TRIzol reagent (Invitrogen), stored at -20°C for up to a week and processed according to manufacturer’s instructions with slight adaptations. In short, samples were brought to RT for 5min, gently mixed with 200µl 100% chloroform for 3 min by repeated inversion and centrifuged at full speed (16.000 g) at 4°C. The aqueous phase was transferred and mixed with an equal volume of 100% isopropanol by inversion, and incubated for 15 min at RT. RNA was pelleted by centrifuging for 15min at full speed (16 000 g, 4°C), washed with freshly prepared 80% ethanol and centrifuged again. All ethanol was carefully removed and dried off on a heated block at 60°C. RNA was resuspended in 10µl of ultrapure water and digested with 2U TurboDNase (Invitrogen) in 30µl for 30min at 37°C. RNA was precipitated with 2.5M LiCl (from MEGAscript® T3 kit, Invitrogen) and 20 µg of glycogen (Roche) at -20°C for 30 min. RNA was pelleted by centrifuging for 15min at full speed (16 000 g, 4°C) and washed with ice cold 80% Ethanol. After air-drying the pellet, RNA was resuspended in ultra-pure water and concentration was measured on a NanoDrop (Thermo Scientific).

### Genomic DNA extraction

Half of the cell pellets obtained from ACME dissociation and Click reaction was used for DNA extraction and stored in TE Buffer (10mM Tris pH 8, 1mM EDTA pH 8, 25mM NaCl) at -20°C for up to a week. After thawing, 100µg/ml Proteinase K was added and the cells were incubated at 50°C until fully digested (15-20 min). Each sample was mixed with 1ml of phenol by inverting and centrifuged for 3 min at full speed (16 000 g) at RT. The supernatant was transferred to a fresh tube mixed with 1ml of phenol/chloroform (1:1 mix) and centrifuged for 3 min at full speed. The supernatant was mixed with 1ml of 100% chloroform and spun down for 3 min at full speed. Again, the supernatant was transferred and precipitated with 100 µl 3M Na-Acetate pH 5.2 and 1ml of ice-cold 100% ethanol. DNA was pelleted by centrifugation for 10 min at full speed (16 000 g, 4°C), washed with ice-cold 70% ethanol and spun down again. After the ethanol was removed, the pellet was air-dried and resuspended in TE buffer. The DNA was then incubated with 100µg/ml RNase A for 15 min at 37°C and precipitated again using 0.1 vols 3M Na-Acetate and 2.5 volumes of ice-cold 100% ethanol. DNA was pelleted by 10 min centrifugation at full speed (16 000 g, 4°C) and washed with 80% ice-cold ethanol. After removal of ethanol and air-drying, the DNA pellet was taken up into ultra-pure water and its concentration was measured on a NanoDrop (Thermo Scientific).

### Typhoon imaging

ACME fixed cell suspensions of EU-labelled animals and controls were split after coupling to Alexa Fluor™ 488 for genomic DNA and total RNA extractions (see above). 0.4 µg of total RNA and genomic DNA were blotted on an Amersham Hybond^TM^ -N^+^ membrane and the Alexa488 fluorescence was read with a Typhoon biomolecular imager (FLA 9000) using a 473 nm laser with the standard LPB filter.

### Animal culture, EU and BrdU co-labeling and imaging in *Exaiptasia pallida* polyps

Aposymbiotic clonal lines of *Exaiptasia diaphana* strain CC7 (Grawunder et al., 2015) were maintained at 25°C in 1.5 l plastic boxes in the dark with filtered sea water at a salinity of ±32‰. Polyps were fed 5 days per week, with a full exchange of water twice a week, and a monthly scraping of the boxes. All experiments were performed on small polyps originating from clonal reproduction collected a few days before experiments to allow for recovery.

Polyps were labelled for two hours in filtered sea water with 2% DMSO, 50mM MgCl_2_ 0.5mM EU and 0.1mM BrdU. After a brief rinse with sea water containing 50mM MgCl_2_, animals were fixed in 1x PBS, 3.7% FA, 0.2% TritonX, 0.5% DMSO for a 30 min at RT, followed by an overnight fixation at 4°C in fresh fixation solution. Washes and Click-reaction and immunostaining for BrdU was performed as for *Nematostella*, described above. Half of the animals were then treated with 200µg/ml RNase A (PureLink™, Invitrogen) in PTx for 15 min at 37°C and washed 5 times in PTx before mounting for confocal imaging.

Fixed whole animals were mounted with ProLong Gold^TM^ (Invitrogen) and imaged on an Olympus FLUOVIEW FV3000 microscope with standard PMT detectors and a 60x silicon-immersion lens. Images of epidermal regions were recorded as z-stacks with laser power and sensor sensitivity kept identical between samples. All images were consistently adjusted for brightness and contrast in Fiji/ImageJ, except for the DNA stain that was adjusted differently between images.

### Culture, EU and BrdU co-labelling and imaging in HEK293T cells

HEK293T cells were thawed and grown for <5 population doublings in petri dishes in DMEM (#41965, Gibco), 10% FBS at 37°C with 5% CO_2_ in a humidity-controlled incubator. For experiments, cells were seeded into poly-D coated glass bottom dishes (Ø 2cm) and grown to 80% confluency.

For labelling, the cells were incubated in DMEM with 0.1mM BrdU and 0.5 mM EU in 2% DMSO for 2h at 37°C in the incubator, washed with DPBS (#14190, Gibco), and fixed for 30min with 4% PFA (#28906, Thermo Scientific). After permeabilizing for 10min with PBTx (1x PBS, 3% BSA, 0.2% TritonX), cells were washed with 1X PBS and clicked to Alexa Fluor™ 488 azide for 30min as per manufacturer’s instructions.

After a brief wash with PTx (PBS, 0.2% TritonX), DNA was hydrolysed by incubation with 2M HCl for 10min at RT and cells were rinsed with PBTx. Blocking was performed in PBTx with 5%NGS (blocking buffer) for 15min and primary anti-BrdU rabbit antibody (PA5-32256, Invitrogen,) in 1:500 in blocking buffer was incubated with the cells for 2h at RT on slow orbital shaking. After three washes with PBTx, secondary goat-anti-rabbit-Alexa568 antibody (A-11011, Invitrogen,) was added 1:1000 along with 1:5000 Hoechst33342 in blocking buffer and cells were kept overnight at 4°C on slow orbital shaking. Secondary antibody was removed by three washes in PBTx.

Half of the samples were incubated with 200µg/ml RNase A (PureLink™, Invitrogen) in PBS for 15 min at 37°C and washed three times with PTx before confocal imaging on an Olympus FLUOVIEW FV3000 microscope with standard PMT detectors. Cells were imaged using a 60x silicon-immersion lens as z-stacks with laser power and sensor sensitivity kept constant. Across all samples, Alexa Fluor signals were consistently adjusted for brightness and contrast in Fiji/ImageJ, except for the Hoechst33342 signal that was adjusted per image.

### Animal culture, EU and BrdU co-labelling and imaging in *Mnemiopsis leidyi* cydippid larvae

Lobate adults (3-5 cm) of wildtype *Mnemiopsis leidyi* were maintained and spawned as previously described [36]. Fertilized eggs were collected and incubated in aerated 1 litre glass beakers for 24 hours until hatching. For labelling assays, cydippids within 24 hours post fertilization were transferred to 12-well plates and incubated for two hours in artificial sea water (ASW) 2% DMSO containing a combination of 0.5 mM EU and 0.1 mM BrdU. Following incubation, animals were fixed in 3.7% formaldehyde with 0.1% Tween-20 in ASW for one hour at RT. Samples were washed three times with PTw (1x PBS, 0.1% Tween-20) and stored at 4°C for up to two days.

EU detection was performed using the Click-iT™ Plus EdU Cell Proliferation Kit for Imaging, Alexa Fluor™ 488 (Invitrogen), which resulted in a brighter EU signal intensity compared to the Click-iT™ Nascent RNA Capture Kit. Samples were blocked in PTw 3% BSA, washed once with 1x PBS + 0.5% TritonX, and once with PTw. The Click-iT reaction was carried out according to the manufacturer’s instructions for 30 min at RT, after which all samples were rinsed with PTw.

BrdU detection was carried out as described for *Nematostella* using the same reagents (see above). Samples were incubated for 2 hours at 4°C with a secondary antibody solution containing goat-anti-rabbit-Alexa568 antibody (Invitrogen; 1:1000) and Hoechst 33342 (1:2000). RNA digestion was performed on half the samples by incubating with RNase A (200 µg/mL in PBS) for 15 min at 37°C.

All samples were mounted in SlowFade Glass Soft-set Antifade Mountant (S36917, Invitrogen) using 100 µm spacers (two layers of Scotch tape) and sealed with CoverGrip Coverslip Sealant (23005, Biotium). Confocal imaging and image processing were performed as described for *Nematostella*.

### Animal culture, EU and BrdU co-labelling and imaging in *Drosophila melanogaster* larval imaginal discs

*Drosophila melanogaster* of the *w1118* strain were cultivated as previously described [37]. Shortly, flies were grown on solid media containing 32.7g dried potato powder, 60g sucrose, 27.3g dry yeast, 7.3g agar, 4.55ml propionic acid and 2g nipagin per litre, at 25°C, 70% humidity, with a 12-hour day/night cycle. Wing imaginal discs were dissected from third instar larvae in Schneider’s Drosophila Medium (CAT: L0207-500).

Dissected wing imaginal discs were incubated in Schneider’s Drosophila Medium containing 0.5mM EU, 0.1mM BrdU, 2% DMSO for 1 hour at RT under rocking. After a rinse with media, the tissue was fixed for 20 minutes in 4% formaldehyde/PBS. Fixed samples were rinsed three times in PTx (1x PBS, 0.1%Triton-X), before blocking for 15 minutes in PBTx (1x PBS, 0.1% Triton-X, 0.5% BSA). After a rinse with PBS the Click-IT™ reaction to Alexa Fluor™ 488 azide was performed for 30 minutes in the dark at RT, according to the manufacturer’s protocol (Click-iT™ Nascent RNA Capture Kit, Invitrogen). Samples were rinsed 5x in PTx before incubation in 2M HCl/ddH_2_O for 30 minutes at RT. After three rinses and four 20-minute washes in PBTx, Anti-BrdU antibody (Invitrogen, PA5-32256, 1:500, in PBTx) was added and incubated overnight at 4°C. The samples were washed thoroughly in PBTx before donkey anti-rabbit-Alexa Fluor 555 antibody was added (A31572, Invitrogen; 1:1000 in PBTx). Following a 2-hours incubation (at RT with rocking), clicked imaginal discs were washed in PBTx as before and stained with DAPI (D9542, Sigma-Aldrich) for 20 minutes and rinsed 5x with PTx afterwards.

Half the samples for each treatment were incubated with 200µg/ml RNase A (PureLink™, Invitrogen in PTx) for 15 minutes at 37°C. Wing discs were then mounted on slides in NPG aqueous mountant (NPG/ glycerol/ phosphate buffer) and imaged by confocal microscopy on an Olympus FLUOVIEW FV3000 confocal microscope with a 60x silicone immersion lens. Imaging setting were kept constant between images. Images were analysed in Fiji/ImageJ where signals were consistently adjusted for brightness and contrast.

### Animal culture, EU and EdU labelling, RNAse treatment and imaging in *Platynereis dumerilii* regenerates

*Platynereis dumerilii* regenerates were produced as previously described [38]. Briefly, specimens were sampled from a laboratory culture (Vienna PIN strain) grown at temperatures between 18°C and 20°C in a 16:8-h light-dark (LD) cycle. Non-reproductive adults of 40-50 segments in size were immobilised in 7,5% MgCl_2_ diluted 1:1 in artificial seawater (ASW). Posterior transverse amputations were performed at segment 20 with surgical scalpel blades (Swann-Morton, Type10, 0101) using a stereoscopic microscope. Animals were transferred to culture boxes with ASW to regenerate up to stage 2 (around 48 hours).

Regenerating animals were incubated in 0.01mM EdU, 0.01mM EU or 0.5mM EU, dissolved in ASW in a glass beaker for 30min at RT. After immobilisation in MgCl_2_/ASW, animals were fixed in 4% Paraformaldehyde (PFA) in 1xPBS with 0.1% Tween-20 (PTw) for 1h shaking at RT. The fixed animals were washed 3×5min in 1xPTx (1xPBS + 0,2% TritonX) and the regenerate including 5 anterior segments were sampled using surgical scalpel blades. The samples were cleared in tissue clearing solution [39] for 10 min at 37°C, washed three times for 5 min in PTx and the click it reaction for EU (Invitrogen, C10329) or EdU (Invitrogen, C10337) was performed following the manufacturer’s guidelines. The samples were washed three times 5 min in PTx and incubated in Hoechst 33342 1:1000 in PTx for 30 min at RT, followed by three times 5 min washing in PBS. Optionally, RNase treatment was performed with 200µg/ml RNaseA (EN0531, Thermo Scientific) in PTx for 15min at 37°C. After three times 5 min washing with PTx, samples were stored in Vectashield antifade mounting medium (Vector Laboratories, H-100-10) o/n at 4°C and mounted on glass slides and with a coverslip. Confocal images were taken on a Zeiss LSM900 confocal microscope with a LD LCI Plan-Apochromat 40x/1.2 Multi-Immersion, WD 0.41 mm lens, using Zeiss Zen software. Images were processed using FIJI/ImageJ for adjustment of contrast, coloration and creation of overlay images.

## Results

### EU strongly labels nuclei of proliferating cells in the sea anemone *Nematostella vectensis*

To study *de novo* RNA transcription in the sea anemone *Nematostella vectensis*, we adapted the Click chemistry-based EU based protocol [22] to visualise and quantify RNA transcripts in fed juveniles polyps (Fig. 1A). Using flow cytometry (Supplementary Fig. 1, Additional file 1 & Methods), we first benchmarked EU at different concentrations (0.5mM, 1mM and 5mM) to determine the optimal working concentration (Supplementary Fig. 2, Additional file 1). We found that 2h incubation with 0.5mM and 1mM resulted in small fractions of EU-labelled cells among all cells (i.e. EU index), whereas at 5mM EU the EU index was significantly decreased (Supplementary Fig. 2A, Additional file 1). We also saw that the fluorescent intensity of EU+ cells decreased with increasing EU concentrations (Supplementary Fig. 2B, Additional file 1). This result indicated that higher EU concentrations may inhibit uptake or integration, and we therefore chose 0.5mM as standard working concentration for EU incubation.

**Fig. 1.**
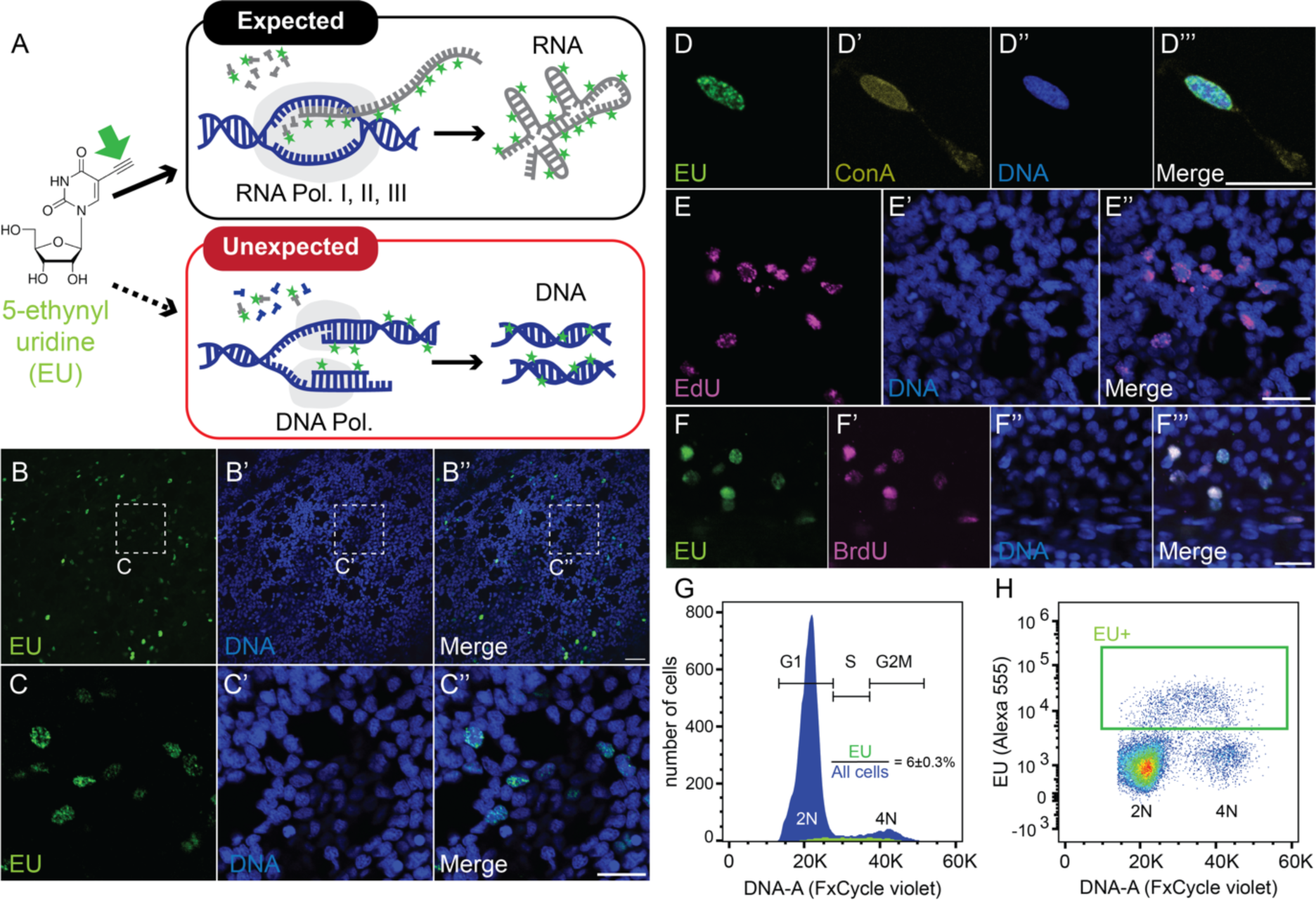
EU labels the nucleus of proliferating cells. **(A)** Schematic overview of EU integration by RNA polymerases into RNA, or alternatively by DNA polymerases into replicating DNA during the S-phase of the cell cycle. Green arrow highlights the ethynyl moiety used to couple a fluorophore by click chemistry. Green stars represent fluorophores. **(B-F’’’)** Single confocal images of epidermal cells labelled for EU (B-D’’’), EdU (E-E’’) or co-labelled for EU and BrdU (F-F’’’) in juvenile *Nematostella* polyps. C-C’’ represent higher magnification of the areas labelled in B-B’’. D-D’’’ represent high magnifications of a labelled nucleus to show nuclear morphology. Note the absence of any labelled nucleolus-like structures. **(G-H)** Flow cytometry histogram (G) of FxCycle Violet DNA stain or density plot (H) of DNA dye (DNA-A) and EU signal intensity (Alexa555) show that EU+ cells range between 2N and 4N DNA content. G1, S and G2M highlight the ranges of DNA content typically found within G1, S or G2/M cell cycle phases of diploid cells. Colour gradient in (H) represents event density from high (red) to low (blue). ±: represents standard error of the mean. Flow cytometry: n=3 replicates. Nuclear stain (blue in B’-C’’ and E’-F’’): Hoechst33342. ConA: Concanavalin A. K: 1.000; Pol.: Polymerase. Scale bars: 20µm (B’’) or 10µm (C’’, D’’, E’’, F’’’).

Using confocal imaging, we found that labelling juvenile *Nematostella* using a 2-hours long pulse of 0.5mM EU resulted in an unexpectedly small subset of cells showing high EU signal (Fig. 1B-C’’). Among these EU+ cells, the fluorescent signal appeared broadly distributed throughout the nuclei (Fig. 1C, D-D’’’), unlike the intense signal in the nucleolus described from EU labelling in other systems [23, 40]. To test if active mRNA transcription is unexpectedly limited to a small fraction of cells in *Nematostella* juveniles, we used immunofluorescence to detect phosphorylated Ser-2 (pSer2) on RNA polymerase II C-terminal repeats that mark active transcription elongation. In contrast to the EU signal, pSer2 was detected in all cells (Supplementary Fig. 3, Additional file 1).

The spatial distribution and relative proportion of EU+ cells were reminiscent of the proliferative cells in fed juveniles labelled with 5-ethynyl 2’-deoxyuridine (EdU), a 2’-deoxynucleoside analogue that integrates into DNA during S-phase [32, 41]. A direct comparison showed that indeed, the EU signal strongly resembles the EdU signal after a 2 hours-long incubation both in appearance and abundance (compare Fig. 1C-C’’ with 1E-E’’). As both EU and EdU contain ethynyl moieties, they cannot be independently click-labelled with two different alkynylated fluorophores. We therefore co-incubated EU with BrdU as alternative to EdU and found both labels widely co-staining the same nuclei throughout the epidermis (Fig. 1F-F’’’). To test if EU+ cells are indeed undergoing DNA replication during S-phase, we quantified their DNA content using flow cytometry (Fig. 1G, H; see Supplementary Fig. 1, Additional file 1, and Methods for gating strategy). We confirmed that the EU+ cells constitute only a small fraction (6±0.3%) of all cells (Fig. 1G) and that their DNA content ranges between 2N and 4N as is typical for S-phase cells (Fig. 1H). Altogether, our results thus indicate that EU consistently labels nuclei of S-phase cells in *Nematostella*.

### EU signal in proliferating cells is insensitive to RNase A, RNase H and the transcriptional inhibitor actinomycin D

In many organisms, DNA replication is accompanied by mRNA transcription during S-phase [42]. Also, DNA primase enzymes generate RNA primers necessary for the initiation of DNA replication of both the lagging and leading strand [43]. We therefore tested if the strong nuclear label of EU is sensitive to RNase A, which degrades single-stranded RNA [44], or to RNase H that degrades RNA/DNA hybrids [45]. We found that neither RNase A nor RNase H treatments had any effect on the intensity of the EU or EdU signal (Fig. 2C-F), supporting that both EU and EdU integrated into DNA in *Nematostella*.

**Fig. 2.**
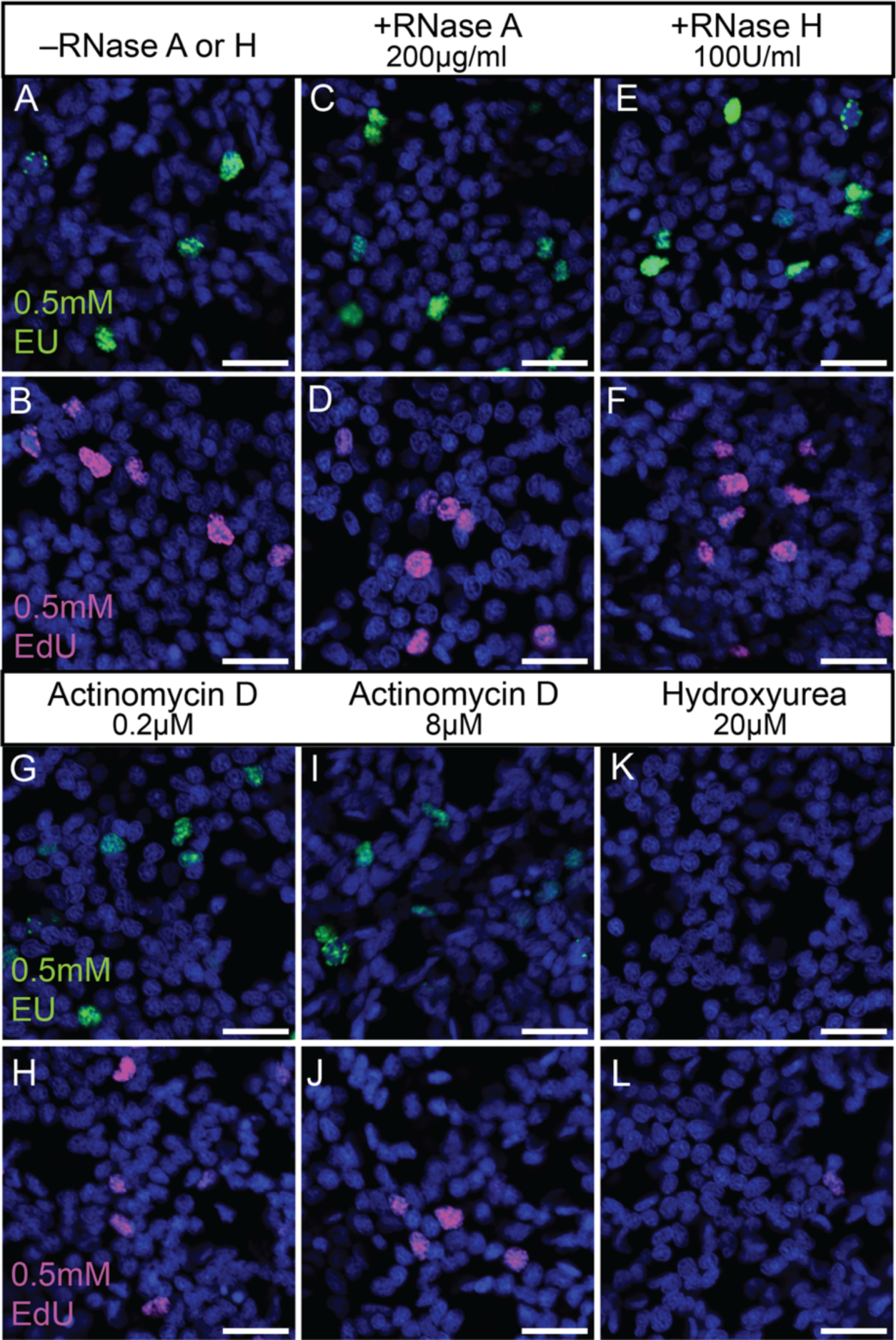
Nuclear EU label in proliferating cells is insensitive to RNase A, RNase H and actinomycin D, but sensitive to hydroxyurea. **(A-L)** Single confocal images of epidermal cells labelled for EU (A, C, E, G, I, K) or EdU (B, D, F, H, J, L) in juvenile *Nematostella* without any treatment (A, B), or treated with RNase A (C, D), RNase H (E, F), the transcriptional inhibitor ActD at 0.2µM (G, H) and 8µM (I, J), or the ribonucleotide reductase (RNR) inhibitor hydroxyurea (K, L). Nuclear stain (blue): Hoechst33342. Scale bars: 10µm.

Furthermore, we reasoned that if EU labelled DNA, the signal should be insensitive to RNA transcription inhibitors such as actinomycin D (ActD), which inhibits RNA synthesis by intercalating into transcriptionally active DNA [46, 47]. ActD has previously been used as inhibitor of RNA transcription to validate that EU integration into RNA is transcription-dependent in cultured cells [22]. In *Nematostella*, we observed that nuclear EU or EdU staining was not affected by low (0.2µM) or high concentrations (8µM) of ActD (Fig. 2G-J). Nuclear EU labelling in proliferative cells of the *Nematostella* juvenile epidermis is thus insensitive to transcription inhibition by ActD and to RNase A and H treatments, leading us to conclude that EU integrates into DNA during S-phase.

The integration of EU into DNA may be explained by a multistep conversion of EU to EdU that involves a reduction of the ribose to 2’-deoxyribose by the RNR enzyme, which is highly conserved across all domains of life. We tested this possibility by using hydroxyurea (HU), a selective inhibitor of RNR that broadly arrests the cell cycle in S-phase, for example in the cnidarian *Hydra* [48, 49], likely by a depletion of the dNTP pool [50, 51]. In previous studies, HU treatment was used in mouse and human cell lines to confirm that EU specifically integrates into RNA during transcription, a typically RNR-independent process [22, 52]. As expected, we found that 20µM HU incubation abolishes EdU labelling (Fig. 2L) consistent with a block of cell proliferation in *Nematostella*. Correspondingly, HU treatment also abolished EU-labelling of nuclei in proliferating cells (Fig. 2K). This further confirms that EU integrates into DNA of proliferating cells, possibly after being converted into EdU by RNR.

Finally, we used a dot blot approach to test if EU is detected in total RNA or genomic DNA of incubated *Nematostella* juveniles (Fig. 3A). We found EU signal in both the RNase A-treated DNA (Fig. 3D-G), sensitive to DNase I (Fig. 3G) as well as DNase I-treated RNA fractions (Fig. 3B), showing that in *Nematostella*, EU integrated not only into DNA, but also into RNA.

**Fig. 3.**
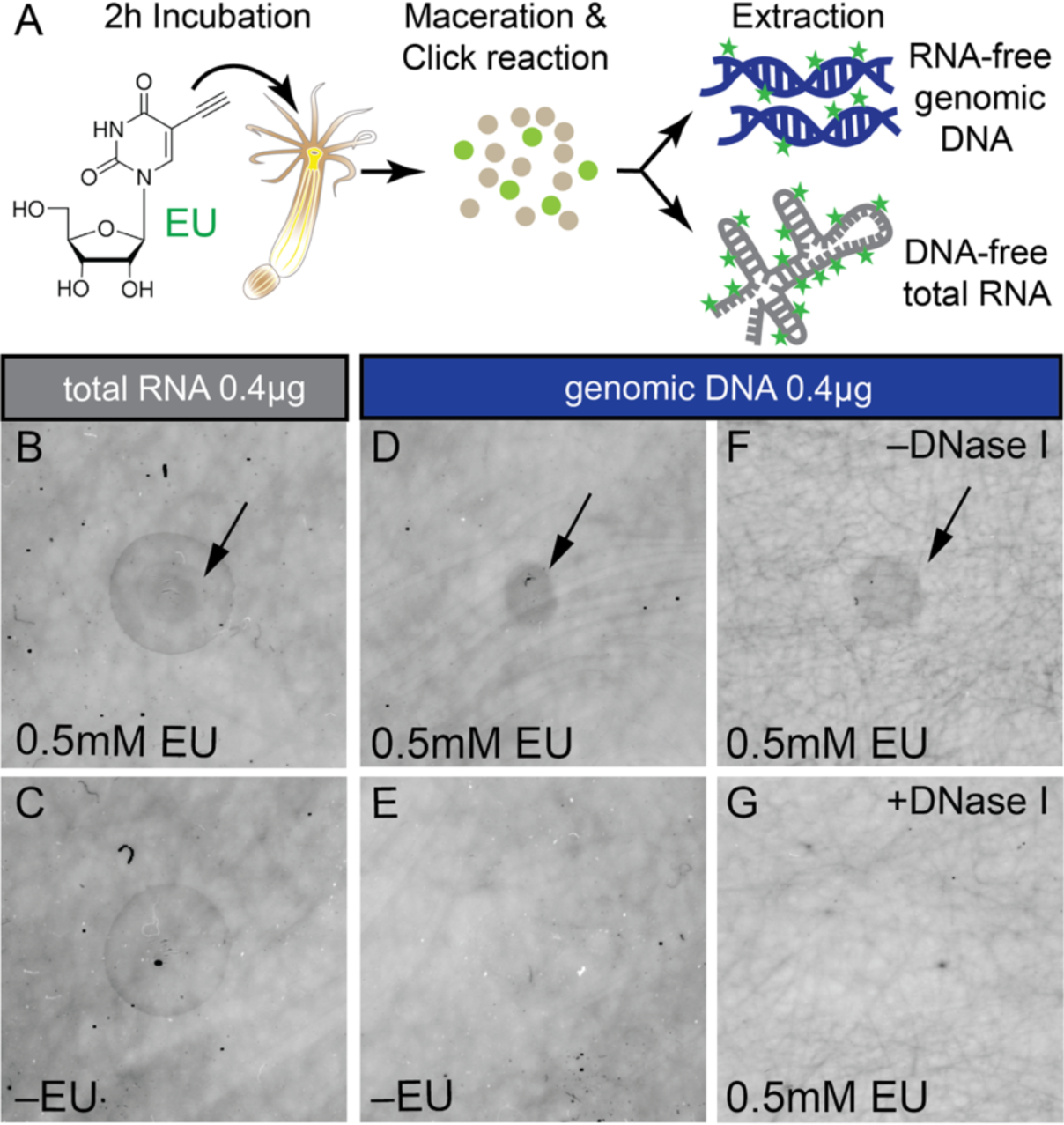
EU is detected in extracted genomic DNA and total RNA. **(A)** Schematic summary of experimental setup. See Methods for details. **(B-G)** Dot blots of DNA-free total RNA (B, C) or RNA-free genomic DNA (D-G) from animals incubated with (B, D, F, G) or without EU (C, E), and treated without (F) or with DNase I (G). Sample volume put on blot corresponded to between 2 and 10µl.

### EU also weakly labels RNA in small, nucleoli-like structures

Finding that EU integrates not only into DNA but also RNA in *Nematostella* led us to revisit confocal imaging of EU-labelled juvenile epidermis using increased laser intensity and detector sensitivities. We reasoned that the integration of EU into RNA should predominantly label nucleoli as sites of high transcriptional activity, as seen in previous cases of unambiguous RNA label [22]. Indeed, we found weak EU signal in small, punctate structures of most nuclei, potentially indicating nucleolar RNA transcription (Fig. 4A-A’’).

**Fig. 4.**
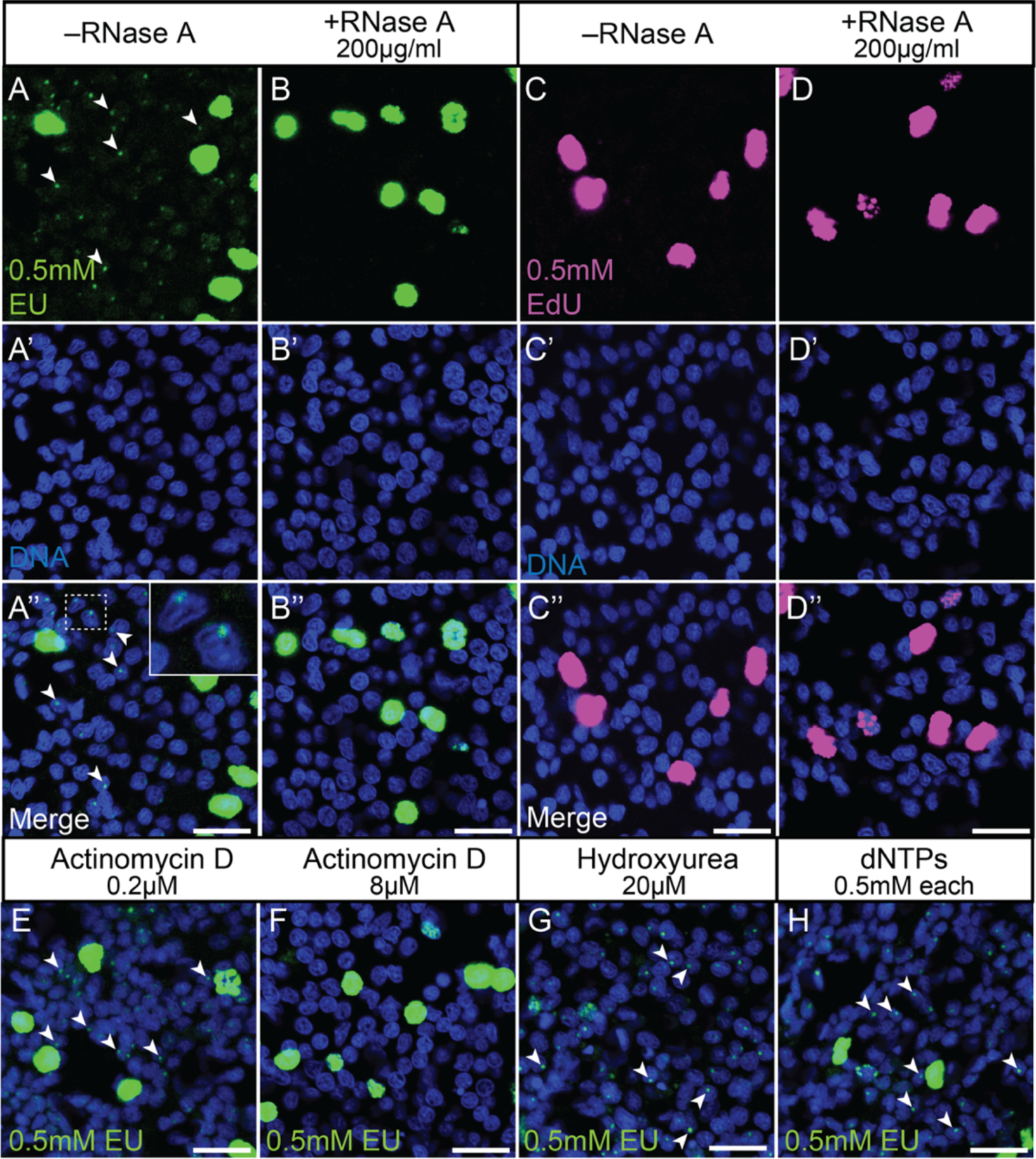
Small, nucleolus-like dots after EU but not EdU labelling are sensitive to RNase A and actinomycin D, but insensitive to hydroxyurea or an excess of dNTPs. Single confocal images of *Nematostella* epidermal cells labelled by EU (A-B’’, E-H) or EdU (C-D’’) and acquired at relatively high laser intensities and detector sensitivities. (**A-B’’**) EU labels small, nucleolus-like dots (arrowheads) that are lost after RNase A treatment. Inlet in A’’ shows higher magnification of the boxed area and depicts labelled nucleolus-like puncta in two nuclei. (**C-D**’’) Using the same detection parameters as in A-B’’, EdU label is specific to nuclei of proliferating cells (C-D’’). **(E-F)** EU label in small nucleolus-like dots (arrowheads) is insensitive to low (E) but abolished by high concentrations (F) of the transcriptional inhibitor ActD. **(G)** Inhibition of ribonucleotide reductase by hydroxyurea leads to reduction of strong nuclear staining, but not of the small nucleolus-like label (arrowheads). **(H)** Adding an excess of dNTPs appears to have no effect on the EU label. Nuclear stain (blue): Hoechst33342. Scale bars: 10µm.

Due to the oversaturation of the fluorescent signal in replicating DNA of proliferative cells, however, we could not determine whether these cells also contain additional EU+ nucleolar puncta. The nucleolar signal was sensitive to 200µg/ml RNase A treatment, while the strong label in proliferating cells remained unaffected (Fig. 4B-B’’). In contrast to EU, incubation with EdU did not label any nucleolus-like structures, indicating that the low EU signal consists indeed of labelled RNA (Fig. 4C-D’’). Aiming to further test this assumption, we studied if the nucleolar-like label was affected by inhibiting transcriptional activity with ActD (Fig. 4E, F). We observed that 0.2µM ActD had no detectable effect on nucleolar EU labelling (Fig. 4E). Instead, 8µM ActD [28] abolished any detectable EU+ puncti (Fig. 4F), indicating that their labelling is dependent on transcriptional activity. Inhibiting RNR by using 20µM hydroxyurea did not affect nucleolar EU staining, while strong nuclear staining previously observed in proliferating cells was lost (Fig. 4G). Altogether, our results support the conclusion that the weak, punctate EU signal indeed corresponds to nascent ribosomal RNA in *Nematostella* epidermal cells. The potential dual integration of EU into RNA and DNA raises the question if supplementation with dNTPs could outcompete EU during DNA but not RNA synthesis. Providing exogenous dNTPs in excess, however, did not result in any major changes in the EU label of the nucleoli or in proliferating cells (Fig. 4H). Altogether, our results suggest that EU is incorporated into both the DNA of proliferating cells and the RNA of transcriptionally active cells in *Nematostella*.

### 5-ethynyl-UTP, 5-ethynyl cytidine and 5-ethynyl adenosine are also not specifically labelling RNA

The dual labelling of DNA and RNA strongly limits the use of EU to visualise and quantify *de novo* RNA transcription levels in *Nematostella*. In search for alternatives, we explored the potential of 5-ethynyl-UTP (EUTP) to specifically label nascent RNA in *Nematostella*, reasoning that it may not act as a direct substrate for RNR and could be directly integrated into RNA [53]. In addition, we tested the labelling characteristics of 5-ethynyl cytidine (EC) and 5-ethynyl adenosine (EA) as potential alternatives. For EUTP and EC, we consistently found a strong, RNase A-insensitive signal in a small subset of nuclei and a weak, RNase A-sensitive signal in small nuclear puncta (Fig. 5A-D’). This strongly indicates that there are no obvious differences in the labelling characteristics between EUTP, EC and EU. In contrast, EA incubation led to broad, RNase A-insensitive labelling of the cytoplasm throughout the epidermis (Fig. 5E, F), especially in longitudinal and circular muscle cells, suggesting that EA gets metabolized and labels ADP/ATP pools (Fig. 5G)[54]. Altogether, we conclude that EUTP, EC or EA are not suitable alternatives to EU for the specific labelling of nascent RNA transcripts in *Nematostella*.

**Fig. 5.**
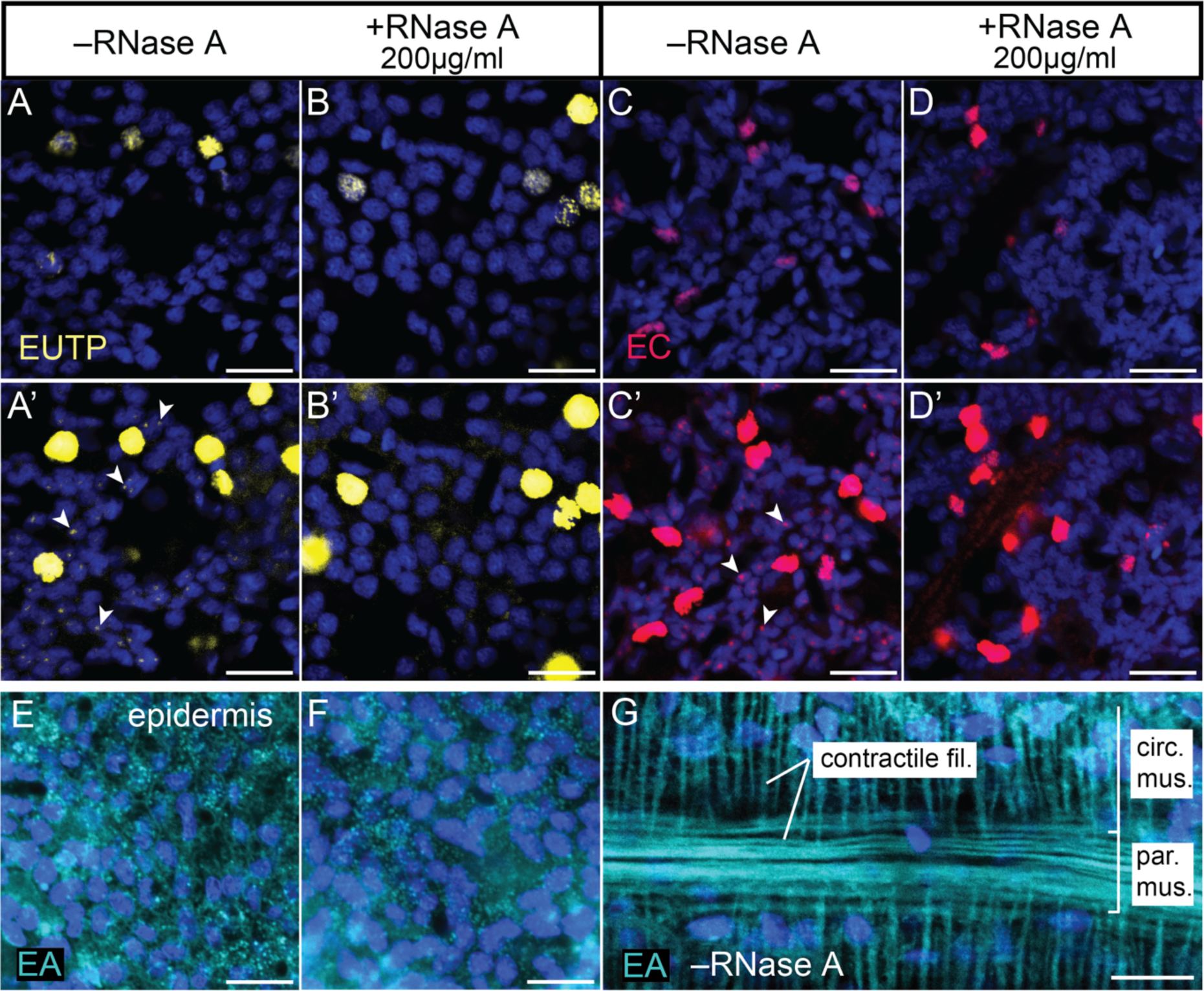
The potential EU alternatives 5-ethynyl uridine triphosphate (EUTP), 5-ethynyl cytidine (EC) and 2-ethynyl adenosine (EA) also integrate into DNA or label other non-RNA structures. Single confocal images (A-F) or confocal imaging stacks (G) of epidermal (A-F) or gastrodermal (G) cells labelled using EUTP (yellow), EC (red) or EA (cyan). (**A-D’**) EUTP (A-B’) and EC (C-D’) are detected in dispersed nuclei of putative proliferating cells, and their signal is RNase A-insensitive (A-D). At high laser intensity and detector sensitivity, both compounds also label small nucleolus-like dots (arrowheads) that are RNase A-sensitive (A’-D’). (**E-G**) EA label is found broadly in epidermis (E) and is insensitive to RNase A degradation (F). EA signal is also prominently found in contractile filaments of muscle cells (G). Nuclear stain (blue): Hoechst33342. circ. mus.: circular muscle; fil.: filaments; par. mus.: parietal muscle. Scale bars: 10µm.

### EU labelling across different systems reveals a diversity of RNA and DNA incorporation

The unexpected incorporation of EU into DNA in *Nematostella vectensis* motivated us to explore whether this phenomenon represents an oddity of *Nematostella*, or occurs more widely in other animal species. We therefore studied EU incorporation in proliferative tissues and cell culture of five additional species, reasoning that an elevated demand for deoxyribonucleotide triphosphates (dNTPs) may favour metabolic repurposing or misincorporation of EU into DNA.

To validate the use of EU for studying transcriptional activity, we examined EU incorporation in two well characterised model systems for cell proliferation: the immortalised human embryonic kidney cell line HEK293T [55] and the growing wing imaginal disc of *Drosophila melanogaster* [56]. In both systems, the EU labelling highlights nucleoli in most cells, is distinct from BrdU and sensitive to RNase-A (Fig. 6F-G’’’, Fig. 6H-I’’’). We thus confirm previous reports that EU specifically labels RNA in HEK293 cells [57] and in *Drosophila* [58].

**Fig. 6.**
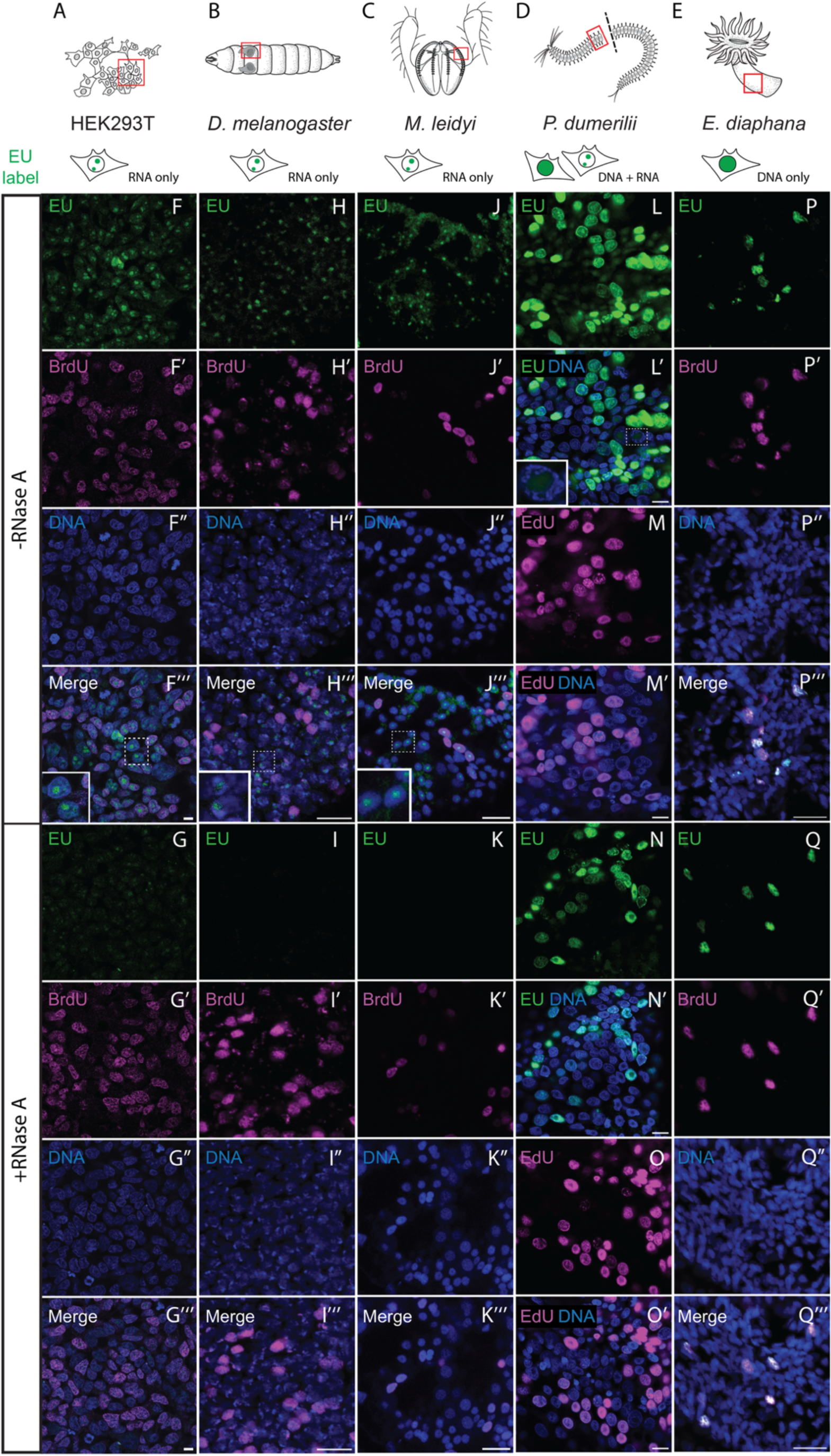
Ethynyl uridine (EU) labelling across diverse model systems reveals variable RNA and DNA incorporation patterns. (A-E) Schematic representation of the examined regions in HEK293T cells, *Drosophila melanogaster, Mnemiopsis leidyi Platynereis dumerilii and Exaiptasia diaphana*, with red boxes indicating imaged tissues and sub-cellular localisation of EU signal summarized below. **(F-G’’’, P-Q’’’)** Co-labelling of EU (green) and BrdU (magenta) in HEK293T cells (F-G’’’), the wing disc of *D. melanogaster* (H-I’’’), the epidermal comb region of *M. leidyi* (J-K’’’) and the epidermal body column of *E. diaphana* (P-Q’’’). **(L-O’)** EU (green) and EdU labelling (magenta) in regenerates of *P. dumerilii*. The samples were either untreated (F-F’’’, H-H’’’, J-J’’’, L-M’, P-P’’’) or treated with 200ug/ml RNase A (G-G’’’, I-I’’’, K-K’’’, N-O’, Q-Q’’’). Inlets in F’’’, H’’’, J’’’ and L’ show higher magnifications of nucleolar EU labelling of nascent RNA. Note that the RNase A-insensitive nuclear EU labelling overlaps with BrdU in *Exaiptasia* (P-Q’’’), consistent with EU integrating into DNA. Also note the nuclear localization and RNase A insensitivity of *Platynereis dumerilii* tail regenerates labelled with either EU (L–L’, N-N’) or EdU (M-M’, O-O’). Samples were untreated (L-N) or treated with RNase A (M-O). Images depict single confocal images (HEK293T, *Platynereis*) or confocal stacks (*Exaiptasia*, *Mnemiopsis*, *Drosophila*). Nuclear stain (blue): Hoechst33342. Scale bars: 10µm.

We then assessed EU incorporation in *Exaiptasia diaphana,* a facultatively symbiotic sea anemone used to study host-symbiont relationship in sea anemones and corals [59-61]. Studying the epidermis of *Exaiptasia* (Fig. 6E), we found that EU co-localises with BrdU to the nuclei of proliferative cells. This EU signal is resistant to RNase A treatment (Fig. 6P–Q’’’), similar our observations in *Nematostella*. As the last common ancestor of *Nematostella* and Exaiptasia goes back to the origin of all sea anamones, the integration of EU into DNA may thus be ancestral to this cnidarian group.

We also explored the incorporation of EU in the ctenophore *Mnemiopsis leidyi* and the polychaete *Platynereis dumerilii*, two marine species used, among other research questions, to study the evolution of nervous systems and regeneration [38, 62-64]. In *Platynereis*, we compared EU and EdU labelling during tail regeneration, a tissue rich in proliferating stem and progenitor cells [65]. Similar to *Nematostella* and *Exaiptasia*, the EU signal was found throughout the nucleus in a subset of cells rather than in punctate, near-ubiquitous nucleoli (Fig. 6L-L’), and thus strongly resembled EdU labelling of proliferating cells (Fig. 6M-M’). The strong nuclear signal of both EU and EdU labelling was RNase A-insensitive (Fig. 6N-O’). In addition, we found RNase-sensitive nucleolar EU signal in some *Platynereis* cells (Fig. 6L’,N’). Together, these results indicate that EU predominantly labels DNA, but also RNA in *Platynereis*.

In the ctenophore *Mnemiopsis leidyi*, we studied epidermal cells around the developing comb rows (Fig.6C) and found that although EU and BrdU co-label some cells, the EU signal is distinctly localising to the nucleoli (Fig. 6J), where no BrdU signal was observed, and is sensitive to RNase A digestion (Fig. 6K’’’). Together, these observations indicate that EU predominantly labels RNA in the *Mnemiopsis* epidermis.

Together, these results demonstrate that EU cannot universally be relied upon as a specific label of nascent RNA. Instead, the specificity to RNA varies markedly across species and tissues, underscoring the need to consider and control for potential DNA integration when using EU.

## Discussion

Combining EU incubation with click chemistry-based fluorophore labelling has promised a quick, easy and commercially accessible method to visualise nascent RNA and transcriptional activity. In this study, contrary to our expectations, we found that EU considerably labelled nuclear DNA of proliferating cells in the sea anemones *Nematostella vectensis* and *Exaiptasia diaphana,* and in the annelid *Platynereis dumerilii*. Our conclusions are based on colocalization between EU and BrdU, on similar staining between EU with EdU, and on the RNase A insensitivity of the strong nuclear EU signal found in proliferative cells. Our finding that inhibition of ribonucleotide reductase enzymes, which salvage ribonucleotides for DNA synthesis [13-15], markedly reduced EU signal in *Nematostella* further consolidates the notion that EU integrates into replicating DNA.

After biochemically isolating separate RNA and genomic DNA fractions from EU-labelled *Nematostella* juvenile polyps, we detected EU not only in DNA but also RNA. Using increased sensitivity in confocal microscopy, we could indeed also observe RNase A-sensitive EU signal in small, nucleolus-like puncta in most epidermal cells of *Nematostella* and in *Platynereis* regenerates. Furthermore, the weak, nucleolar EU signal was inhibited by high concentrations of the transcriptional inhibitor actinomycin D in *Nematostella*.

Together, our data suggest that EU, as well as the alternatives EUTP and EC, integrate predominantly into DNA and to a lesser extent into RNA in *Nematostella.* The strong nuclear EU labelling found in *Nematostella*, *Exaiptasia* and *Platynereis* corresponds to replicated DNA. This result is compatible with a scenario where RNR is part of a metabolic network salvaging EU into EdUTP. In contrast, the predominant integration of EU into nascent RNA was found in human HEK293T cells, the *Drosophila* wing disc and the epidermis of the ctenophore *Mnemiopsis leidyi*.

We demonstrate the limitations of using EU-click labelling to visualise and quantify nascent RNA in *Nematostella*, *Exaiptasia* and *Platynereis* via confocal imaging, flow cytometry or fluorescence-assisted cell sorting (FACS). Likely, EU and other labelled nucleosides remain useful for RNA sequencing approaches, such as 4sU-Seq (i.e. 4-thiouridine sequencing)[66, 67] or GRO-Seq (Global Run-On Sequencing)[68], as EU-labelled DNA is removed during the sample and library preparation process. It should, however, be noted that high integration rates of EU into DNA would likely result in low labelling yields of EU into nascent RNA.

To our knowledge, we provide the first report that EU labels both RNA and DNA in any animal species or tissues. Given that the dual labelling has been observed for other labelled nucleosides or nucleotides (see also introduction) [17-19], our study reinforces the doubts on their indiscriminate use to specifically label RNA, which is rarely verified in direct and systematic ways. Strongly proliferating tissues may be especially prone to dual labelling as their high demands in dNTPs may favour channelling EU towards DNA replication by conversion to EdUTP. While thoroughly screening the recent literature, we found recurring reports of EU strongly labelling the whole nucleus – rather than punctate-like nucleoli – in a subset of cells in highly proliferative tissues. We found examples of EU labelling patterns reminiscent of DNA labelling by EdU or BrdU during *Nematostella* regeneration [28], in the larval zebrafish brain [69], the mouse intestinal crypt cells [22, 52], spermaries [70], granulosa cells [71], embryonic cortex [72] and cerebellum [22, 73]. Our report shows that the labelling behaviour of EU is difficult to predict across animals and tissues, and emphasises the requirement for robust controls. Commonly, the specificity of RNA labelling is validated by using transcriptional inhibitors such as ActD [22, 73], or α-Amanitin [74, 75]. Previous reports show however that the resulting transcriptional stress may induce DNA damage and lead to cell cycle inhibition [76]. Transcriptional inhibitors as the sole validation of RNA specificity may thus be insufficient as any loss of EU signal after using ActD or α-Amanitin can also result from inhibiting DNA replication. While ActD was effective in *Nematostella* to discriminate between EU-labelled DNA and RNA, its use might be more problematic in other systems.

Under the premise that RNA is not a substrate of DNases, another problematic validation assay commonly involves showing that the EU label is insensitive to DNase I digestion [77]. This assay does not directly address whether incorporation into RNA and is also prone to fixation artefacts as formaldehyde-based fixation leads to stable, covalent bonds between DNA and proteins that resist DNase I treatment [78]. This phenomenon is obvious in BrdU detection assays, where DNase I digestion of the label-containing DNA is used to enhance antibody accessibility without significantly compromising the BrdU signal [79].

So far, the RNA specificity of EU has been validated most extensively in *in vitro* systems (e.g. NIH 3T3 or HeLa cells), assuming that the results are transferrable to more complex animal systems, such as mice [22, 52]. While we cannot conclusively evaluate the validity of this assumption, our findings underscore the importance of standardized controls to validate the specificity of EU or other nucleotide-based RNA labelling methods in all experimentally relevant contexts. Based on our experience and observations, we found that the following EU-labelling characteristics are indicative of DNA integration:

- Broad labelling of the nucleus without clear nucleolar signal
- Condensed chromosome or metaphase plane labelling in mitotic cells
- Nuclear labelling of a smaller subset of cells in proliferative tissues
- Insensitivity to RNase A treatment
- Insensitivity to transcriptional inhibitors (e.g. ActD)

The occurrence of one or several of these features should prompt further validation experiments. We propose the following good practice guidelines when using EU to visualise nascent RNAs:

- **Validate RNA specificity of EU labelling by RNase A or T1 treatment.** EU signal originating from the integration into RNA should be sensitive to RNase A or T1 treatment. In contrast, EU-labelled DNA will not be removed by RNase A or T1 treatment and may also be insensitive to DNase I treatment in fixed tissues (see above).
- **Investigate high variation of EU labelling in proliferative tissues or in cell culture by direct comparison with EdU or BrdU labelling.** A direct comparison between cells labelled by EU and EdU or BrdU will help revealing differences or similarities in labelling frequency, intensity and location. We propose co-labelling of EU and BrdU on the same tissue as gold standard to test if EU integrates into replicating DNA of proliferative cells.
- **Consider potential cell cycle off-target effects when using transcriptional inhibitors to control for RNA specificity.** Inhibition of transcription often causes cellular stress that can alter the cell cycle, affect DNA synthesis rate or lead to cell death. Any resulting loss of EU signal may therefore be caused by decreased transcription or blocked proliferation.

EU and other nucleoside-based nascent RNA labelling methods have significant limitations due to their potential to be metabolically converted and incorporated into DNA. Our study shows striking differences in EU incorporation patterns across organisms, labelling RNA specifically in some, while predominantly integrating into DNA in others. The variability in the fate of nucleotide analogues likely stems from species-specific differences in nucleoside transport, differences in the activity of salvage pathways or in the specificity of polymerases. Beyond presenting a technical obstacle for studying transcription, the differences in RNA/DNA incorporation of EU also highlight an intriguing diversity in how organisms use and process nucleotides. Understanding the nuances of nucleotide metabolism across the tree of life will therefore be essential for developing more specific RNA labelling tools, particularly for species and contexts in which conventional nucleoside analogues like EU predominantly label DNA.

## Conclusions

Our study reveals that the nascent RNA labelling compound 5-ethynyl uridine (EU) predominantly integrates into DNA rather than RNA in the sea anemones *N. vectensis*, *E. diaphana* and the polychaete *P. dumerilii*. In contrast, EU specifically labels RNA in HEK293T cells, the wing disc in the *fly D. melanogaster* and in epidermal cells of the ctenophore *M. leidyi*. These contrasting findings reveal critical limitations in the use of EU for studying transcription in certain animal models and underscore the need to experimentally address DNA incorporation in unvalidated biological contexts, particularly in highly proliferative tissues. More broadly, the observed variation in EU incorporation patterns also point to underlying species-specific differences in nucleotide metabolism. Understanding these differences will be essential for developing more reliable RNA labelling tools tailored to diverse model organisms.

## List of abbreviations

2N / 4N: DNA content of diploid cells before (2N) and after replication (4N)
4sU-Seq: 4-Thiouridine Sequencing
ANOVA: Analysis of Variance
ASW: Artificial Sea Water
ActD: Actinomycin D
ACME: Acetic acid/Methanol (maceration protocol)
Alexa488: Alexa Fluor™ 488 fluorophore
Alexa555: Alexa Fluor™ 555 fluorophore
Alexa568: Alexa Fluor™ 568 fluorophore
Ara-C: Cytosine Arabinoside (Cytarabine)
BSA: Bovine Serum Albumin
BrdU: 5-Bromo-2’-deoxyuridine
CAT: Catalog number
CC7: Clonal strain designation of *Exaiptasia diaphana*
ConA: Concanavalin A
circ. mus.: Circular muscle
DAPI: 4′,6-diamidino-2-phenylindole (DNA stain)
ddH_2_O: Double-distilled Water
dNDP: 2’-deoxy-Nucleoside Diphosphate
dNTPs: Deoxynucleoside Triphosphates
DMEM: Dulbecco’s Modified Eagle Medium
DMSO: Dimethyl Sulfoxide
DNA: Deoxyribonucleic Acid
DNA-A /H: DNA signal area / height (flow cytometry)
DNase I: Deoxyribonuclease I
DPBS: Dulbecco’s Phosphate-Buffered Saline
EA: 2-Ethynyl Adenosine
EC: 5-Ethynyl Cytidine
EDTA: Ethylenediaminetetraacetic Acid
EU: 5-Ethynyl Uridine
EU+: Cells labeled with 5-Ethynyl Uridine
EUTP: 5-Ethynyl Uridine Triphosphate
EdU: 5-Ethynyl-2’-deoxyuridine
EdUTP: 5-Ethynyl-2’-deoxyuridine Triphosphate
EdU+: Cells labeled with 5-Ethynyl-2’-deoxyuridine
F-ara-A: 9-β-D-arabinofuranosyl-2-fluoroadenine
FA: Formaldehyde
FACS: Fluorescence-Assisted Cell Sorting
FBS: Fetal Bovine Serum
Fil.: Filaments
FSC-A /H: Forward Scatter Area / Height
5-FU: 5-Fluorouracil
5-FUTP: 5-Fluorouracil Triphosphate
5-FdUTP: 5-Fluoro-2’-deoxyuridine-5’-triphosphate
g: Relative centrifugal force (centrifugation)
G1: Gap 1 phase of the cell cycle
G2M: Gap 2 phase and Mitosis phase of the cell cycle
GRO-Seq: Global Run-On Sequencing
3H: Tritium (Hydrogen isotope)
HEK293T: Human Embryonic Kidney 293T (cell line)
HSD: Honestly Significant Difference (Tukey’s Post hoc test)
HU: Hydroxyurea
K: Kilo (1000)
Milli-Q: Ultra-purified water
mRNA: Messenger RNA
NDP: Nucleoside Diphosphate
NGS: Normal Goat Serum
NIH 3T3: National Institutes of Health (NIH) 3T3 mouse fibroblast cell line
NPG: n-Propyl gallate
NTP: Nucleoside Triphosphate
n.s.: Not significant (statistics)
par. mus.: Parietal muscle
PBS: Phosphate-Buffered Saline
PBTx: PBS with TritonX and BSA
PFA: Paraformaldehyde
PMT: Photomultiplier Tube
PTw: PBS with Tween-20
PTx: PBS with TritonX
Pol.: Polymerase
RNA: Ribonucleic Acid
RNA Pol II: RNA Polymerase II
RNR: Ribonucleotide Reductase
RNase A / H: Ribonuclease A / H
RT: Room Temperature
S-phase: DNA synthesis phase of the cell cycle (DNA replication)
SSC-A / W: Side Scatter Area / Width
TE Buffer: Tris-EDTA Buffer
TRIzol: Reagent for RNA extraction
TritonX: Triton X-100
U: Units (enzyme concentration)
µm / µl: Micrometer / liter

## Declarations

### Ethics approval and consent to participate

Not applicable

### Consent for publication

Not applicable

### Availability of data and materials

All data generated and/or analysed during this study are included in this published article and its supplementary information file (Additional file 1). Raw image files are available from the corresponding author on reasonable request.

### Author contributions

MAK, KG and PRHS designed the study and interpreted the results. MAK performed all labelling assays, detection of RNA/DNA on blots, flow cytometry and confocal microscopy for *N. vectensis* and *E. pallida.* KG and MC performed EU/BrdU co-labelling and confocal imaging on HEK293T cells. KLPD conducted tissue dissections, EU/BrdU co-labelling and confocal imaging for *D. melanogaster*. RS performed co-labelling of EU/BrdU and confocal microscopy of *M. leidyi* cydippids. LA bisected, labelled and imaged *P. dumerilii* regenerates with EU and EdU. MAK, KG and PRHS wrote the manuscript with contributions from KLPD, RS, LA, MC, FR, PB and FO’F. All authors read and approved the final manuscript.

## Acknowledgements

We thank Simon Henriet for support and materials for the membrane blotting experiments, Eudald Pascual-Carreras, Jørn Skavland and Brith Bergum for help and support with flow cytometry, Andreas Midlang for help with confocal imaging, Annika Guse for sharing Aiptasia strains, and Eilen Myrvold and Brandon Mellin (*Nematostella*, Aiptasia), Alexandre Jan (*Mnemiopsis*), Sandra Ninzima (*Drosophila*) and Andrij Belokourov, Maragyta Borysova, and Netsaneh Getachew (*Platynereis)* for animal husbandry and maintenance. We are grateful to all members of S12 for critical discussion. The flow cytometry was performed at the Flow & Mass Cytometry Core Facility, Department of Clinical Science, University of Bergen. Except for samples from *Platynereis dumerilii*, all confocal imaging was performed at the Michael Sars Centre and the BIO Department at UiB.

