## Supplementary figures 1-4 for "The nascent RNA labelling compound 5-ethynyl uridine (EU) integrates into DNA in some animals"

### Supporting Information

### contains 4 Supplementary Figures S1-S4

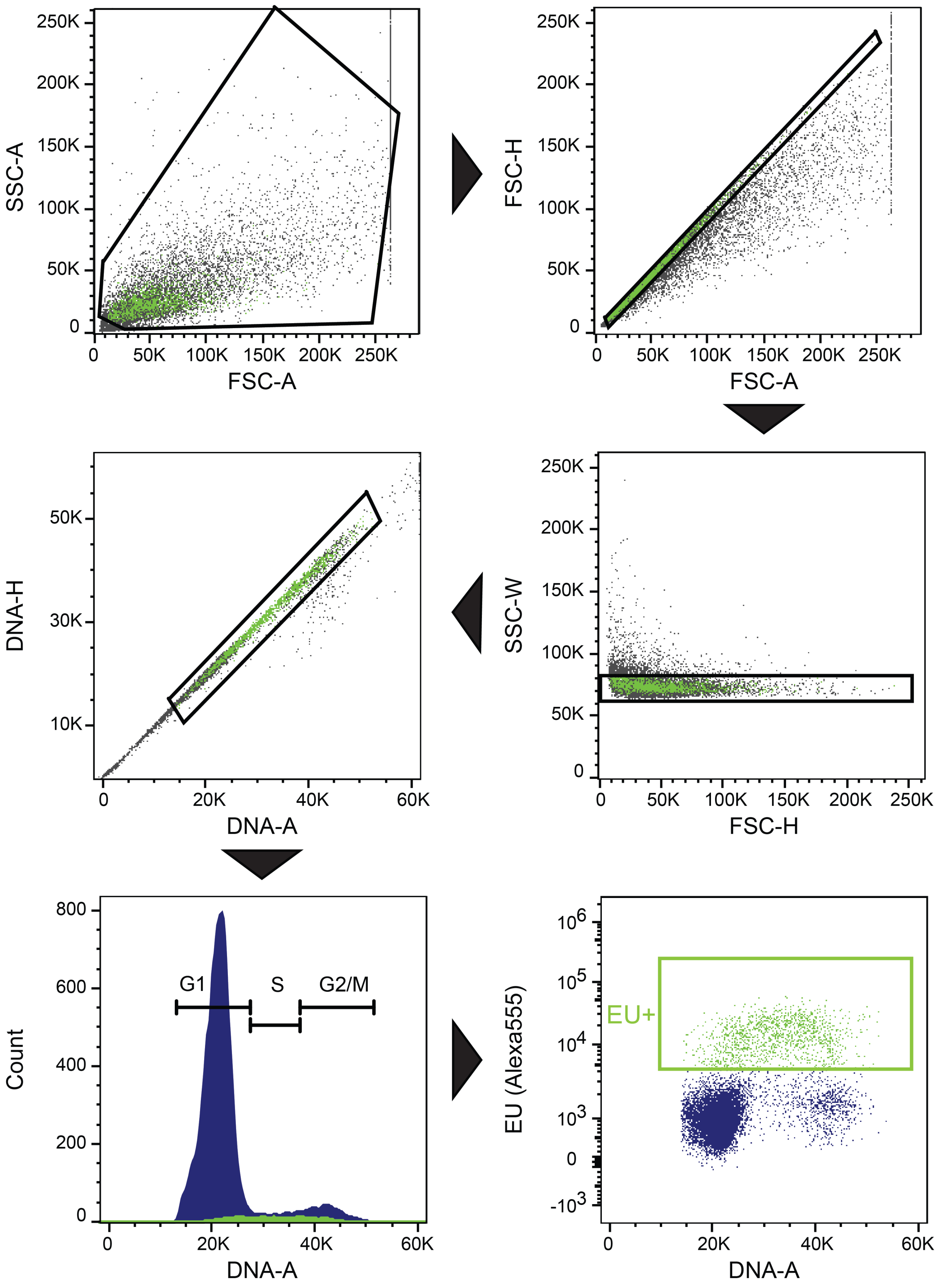
SSC-A
FSC-A
FSC-H
FSC-A
SSC-W
FSC-H
DNA-H
DNA-A
Count
DNA-A
G1
S
G2/M
EU (Alexa555)
DNA-A
EU+

**Supplementary Fig. 1. Gating strategy to detect EU signal (Alexa 555) using flow cytometry.** For gating cells, very small events were excluded based on size and granularity in the FSC-A/SSC-A gate, followed by gating for linear relation between FSC-A and FSC-H to remove cell aggregates. We limited the analysis to events with low side scattering by gating in FSC-H/SSC-W and plotted the resulting population based on a linear corelation in FxCycleViolet dye intensity of area and height (DNA-A/DNA-H) within a range that corresponds to 2N to 4N DNA content. This gate contains cells within the cell cycle and constitutes the base population for the calculation of the EU positive cells. Based on the fluorescence signal of DMSO controls in the DNA-A / Alexa555-A plot, we set a threshold gate above which cells were considered EU positive. The EU+ population was backgated and indicated as green overlay in the parental gates.

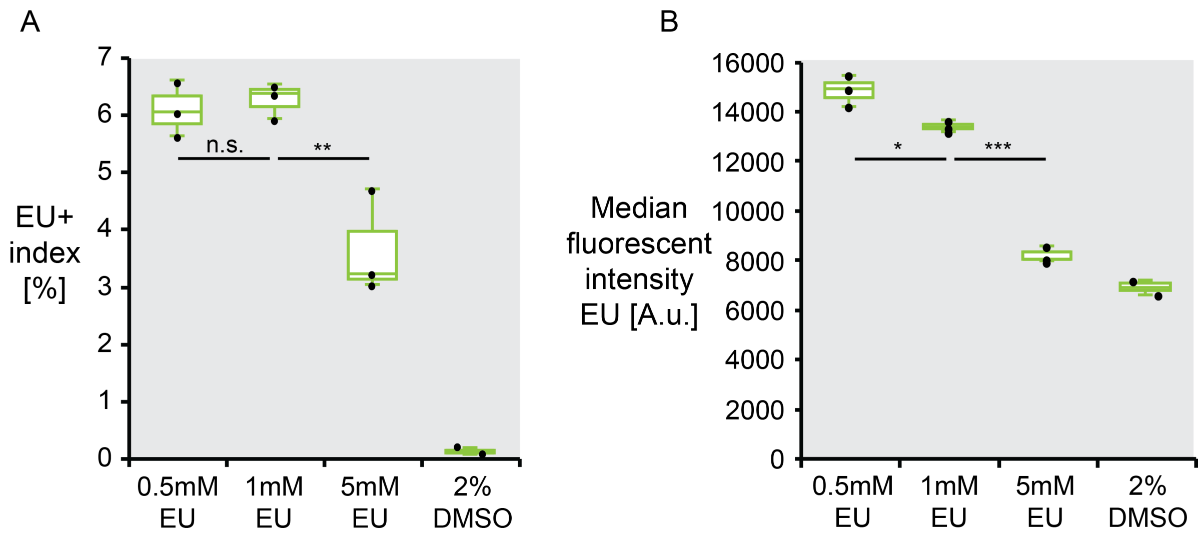


**Supplementary Fig. 2. Fraction of EU positive cells and their median fluorescence after 2 hours of labelling with different EU concentrations.** Box plots depicting **(A)** the fraction and **(B)** the median fluorescent signal intensity of EU+ cells at different concentrations of EU and at 2% DMSO. Box plots indicate median (middle line), 25th, 75th percentile (box) and 5th and 95th percentile (whiskers). Dots show single data points. Differences between concentrations were calculated using One-way ANOVA with Tukey's HSD post hoc test for pairwise comparisons. n.s.: not significant, *: $p > 0.05$, **: $p > 0.01$, ***: $p > 0.001$.

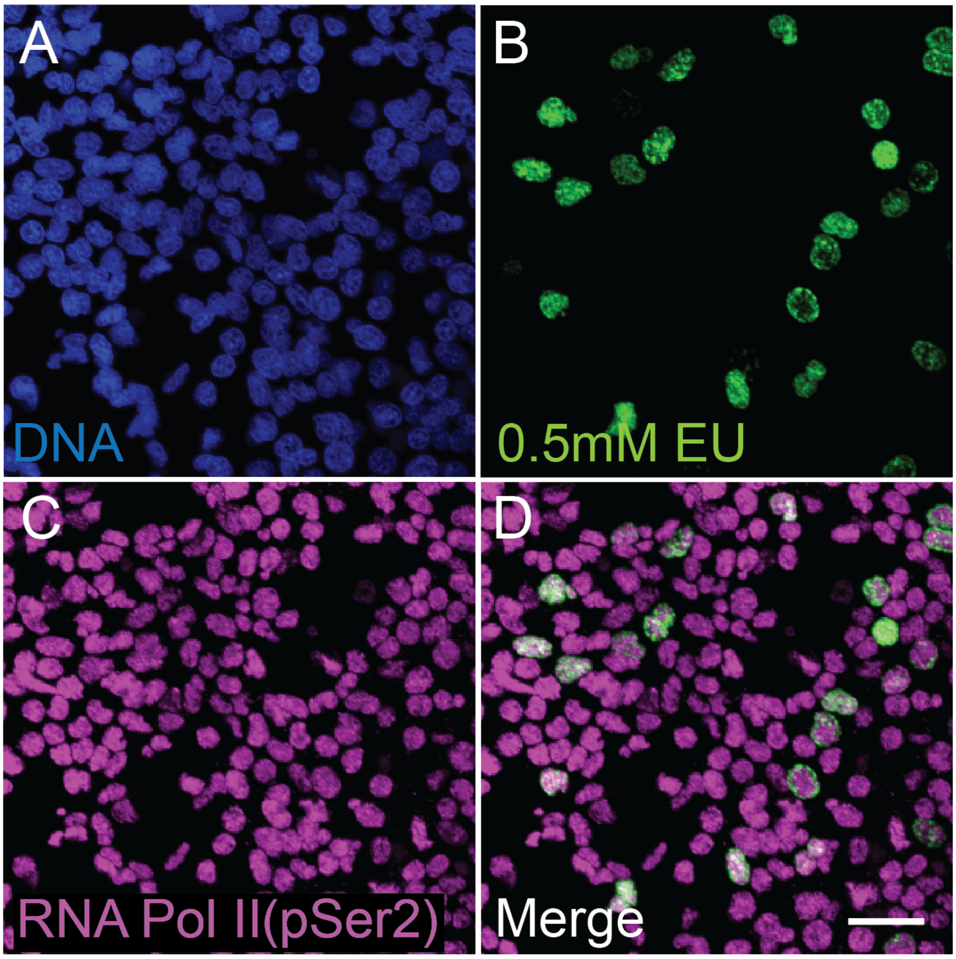


**Supplementary Fig. 3. EU labels a small subset of transcriptionally active cells. (A-D)** Confocal imaging stack of epidermal cells shows that EU (B, D; 2h pulse) labels only a fraction of cells detected by an antibody against phospho-Ser2 of the C-terminal domain of RNA polymerase II (C, D). Nuclear stain (blue): Hoechst33342. Scale bar: 10$\mu$m.

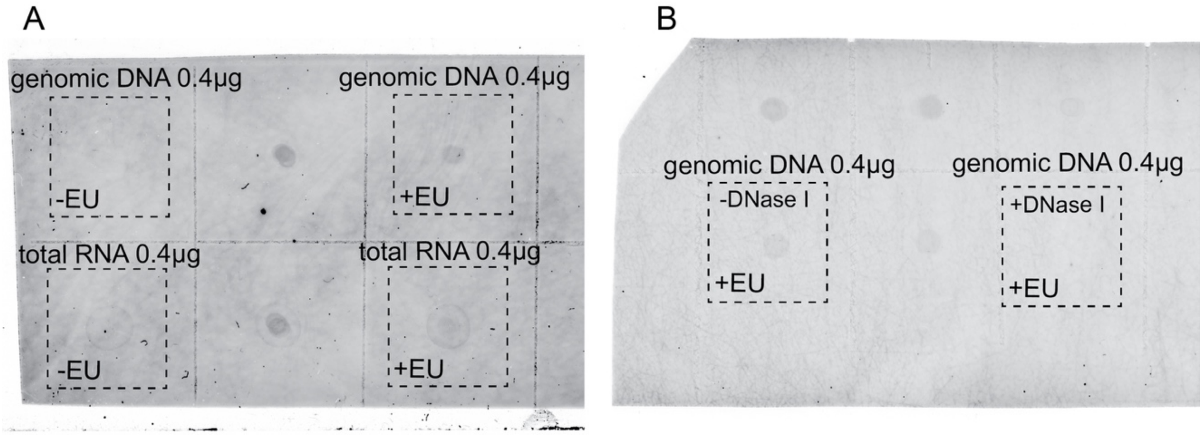


**Supplementary Fig. 4. EU signal in extracted genomic DNA and total RNA.** Dot blots of DNA-free total RNA and RNA-free genomic DNA from animals incubated with EU and without EU **(A)**, as well as with and without DNase I treatment **(B)**. Dashed lines indicate the cropped images shown in Figure 3B-G. Sample volume put on blot corresponded to between 2 and 10$\mu$l to achieve 0.4$\mu$g loading per spot. Relevant sections of the original blots depicted in Figure 3, see Methods for experimental details.
